## supplementary material for "Optogenetic control of a GEF of RhoA uncovers a signaling switch from retraction to protrusion"

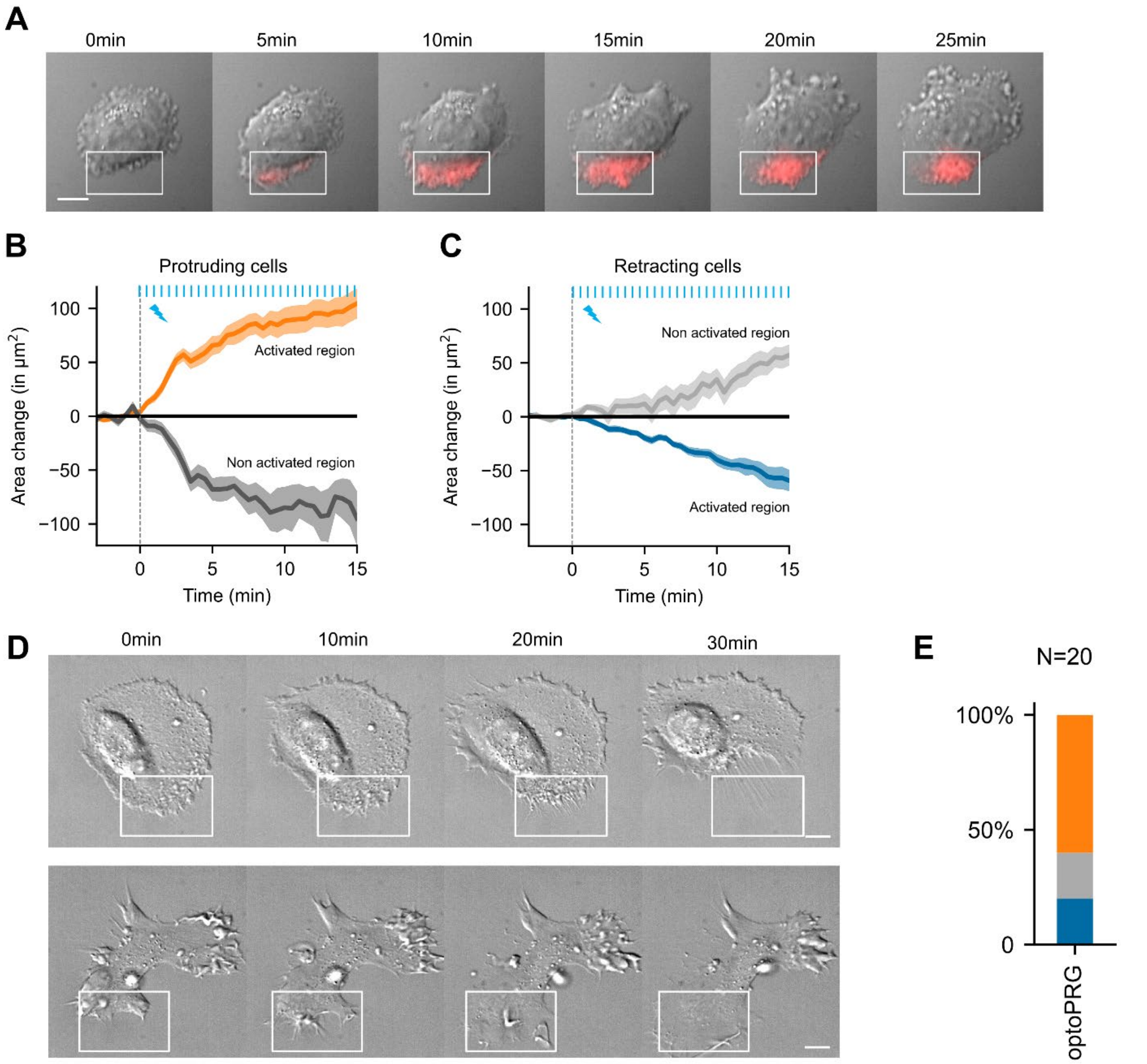


**Supplementary figure 1: (A)** Representative example of a mixed phenotype: here we see first protrusions (from 5 to 10 minutes) and then retraction with blebby structures forming at the opposite side of the cell. Scale bar: 10 µm. White squares: area of optogenetic activation. Red color: RFPt channel (optogenetic tool). **(B, C)** Evolution of the cell areas in the activated region and on the opposite side of the cell for protruding (B) and retracting cells (C). **(D)** Representative Hela cells doing retraction (on the top) and protrusion (on the bottom) upon optogenetic activation. Scale bars: 10 µm. **(E)** Proportion of Hela cells doing a protrusion (orange), retraction (blue) or a mixed phenotype (gray) upon optogenetic activation.

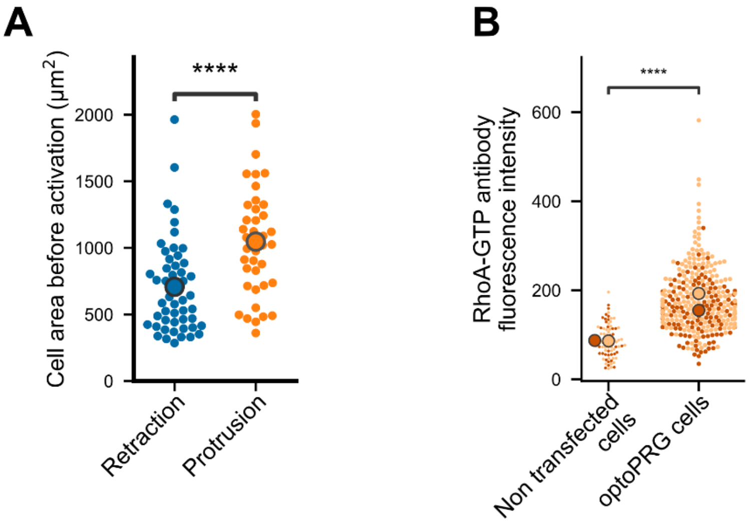


**Supplementary figure 2: (A)** Cell area before activation for retracting (blue) and protruding (orange) cells. **(B)** Quantification of RhoA basal activity in control and optoPRG overexpressing cells, using antibody staining against RhoA-GTP after fixation. Each dot is a single cell, and the two colors correspond to two independent experiments. **** $P<0.0001$ (Mann-Whitney)

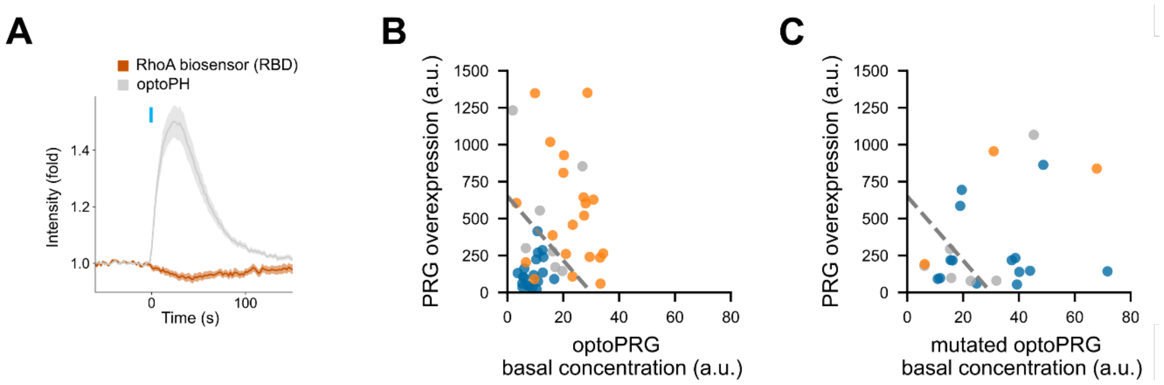


**Supplementary figure 3: (A)** Mean ± sem of fold recruitment of optoPH and RBD biosensor evolution over 26 cells. Light pulse is shown with the blue bar. **(B) (C)** Overexpression of the non-recruitable PRG together with the optogenetic tools. Control experiment **(B)** with recruitment of optoPRG and overexpressed non recruitable PRG, and **(C)** same experiment with the mutated PH. Dashed line separates retracting and protruding phenotypes in control experiment.

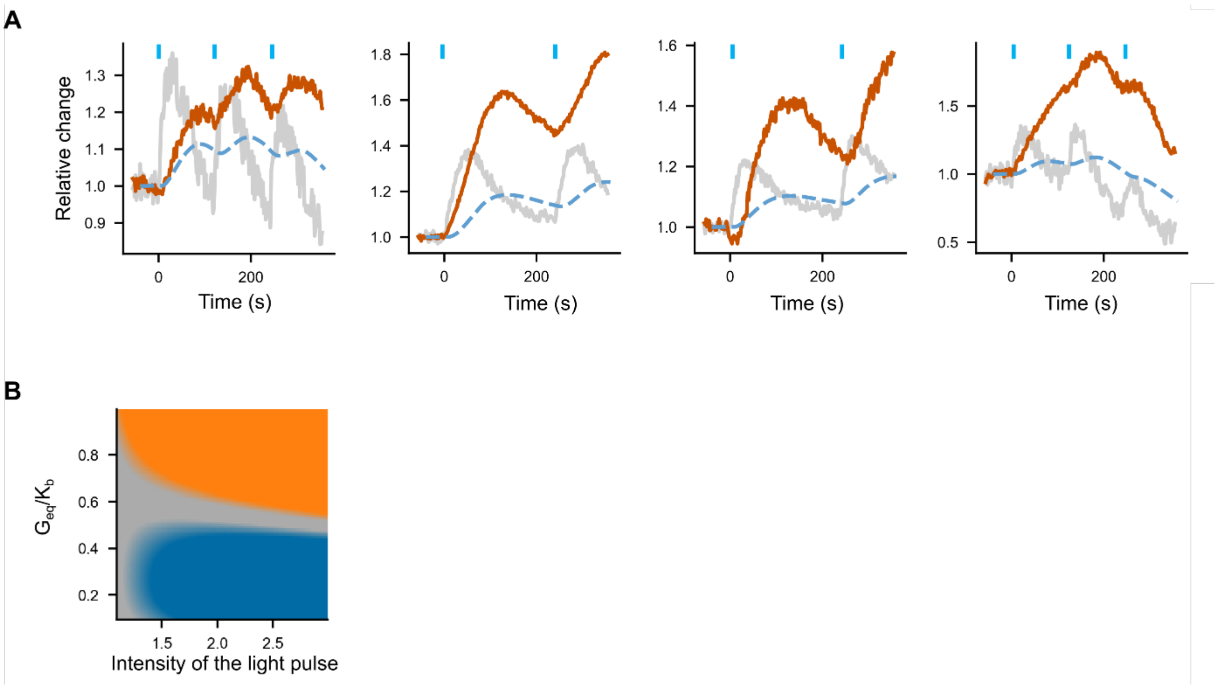


**Supplementary figure 4: (A)** Some biosensor dynamics, shown here, are not well recapitulated by the model. Dotted blue line: fitted curve, with gray line $g(t)$ taken as input (optoPRG recruitment). Red line: RBD biosensor. Chosen value for Geq/Kb is 0.001 for all these plots. **(B)** Phenotype in function of the pulses intensity and Geq/Kb. Protruding (orange), retracting (blue) or mixed (grey) phenotypes are represented in function of the free parameter Geq/Kb (y-axis) and the intensity of the optogenetic pulses, represented by the fold increase on the x axis. There is no way to change from retraction to protrusion or vice versa by changing only the intensity of the light pulse (which would equivalent to moving along a line parallel to the x-axis).
