## supplementary note for "Optogenetic control of a GEF of RhoA uncovers a signaling switch from retraction to protrusion"

### An effective model recapitulates reaction kinetics and enables a control of both phenotypes in the same cell

Given the complexity of all interactions happening, we sought for a minimal model that would capture the simple dependence of the phenotype with optoPRG concentration, as well as the biosensors dynamics.

Let us summarize the different facts that are essential to the model:

- Two opposite polarization phenotypes can be triggered by the recruitment of the same DH-PH domain of PRG at the membrane, in the same cell line, could potentially mean in the same biochemical environment if overexpression of PRG is not affecting gene expression.
- Phenotype can be predicted by the absolute concentration of the DH-PH domain of PRG before activation independently of the recruitment. This means that overexpression has an influence on cell state, defined as protein basal activities or concentrations before activation.
- RhoA, the direct effector of PRG, is activated in the retracting phenotype, but its activation is delayed in the protruding one.
- PH domain binding to RhoA-GTP is required for protruding phenotype but not sufficient, and it is acting as an inhibitor of RhoA activity.
- the DH-PH domain of PRG activates Cdc42, which is required for protruding phenotype.

To keep it as simple as possible, we considered that the resulting phenotype was the result of a competition between the relative amount of RhoA-GTP and Cdc42-GTP after 2 minutes, roughly the time when phenotypic changes start to be seen. Reactions are modeled as mass action kinetics happening at the area of activation - even for enzymatic reactions with the GEF, which is justified if the reaction is happening far from saturation and seems to be the case for GEFs of RhoA at least in another example (Kamps et al., 2020). We considered the optogenetic recruitment as a variation of the total amount of PRG in this area, noted $G_{tot}$. As described above, three interactions of PRG will be considered: the interaction with RhoA-GDP inducing a switch to RhoA-GTP, the interaction with Cdc42-GDP inducing a switch to Cdc42-GTP, and the interaction of the PH domain with RhoA-GTP. This last reaction gives rise to a complex (noted *GR*) that is inhibiting RhoA activity, but doesn't prevent GEF activity, as shown in (Chen et al., 2010).

These assumptions lead to three following reactions:

(a) $\mathrm{RhoA^{GDP}} \underset{\mathrm{k_2}}{\overset{\mathrm{k_1 \cdot G_{tot}}}{\rightleftharpoons}} \mathrm{RhoA^{GTP}}$, with typical concentration $K_a = \frac{k_2}{k_1}$

(b) $\mathrm{RhoA^{GTP}} + \mathrm{G} \underset{\mathrm{k_4}}{\overset{\mathrm{k_3}}{\rightleftharpoons}} \mathrm{GR}$, with typical concentration $K_b = \frac{k_4}{k_3}$

(c) $\mathrm{Cdc42^{GDP}} \underset{\mathrm{k_6}}{\overset{\mathrm{k_5 \cdot G_{tot}}}{\rightleftharpoons}} \mathrm{Cdc42^{GTP}}$, with typical concentration $K_c = \frac{k_6}{k_5}$

In the following, we call $R$ and $C$ the concentrations of $\mathrm{RhoA^{GTP}}$ and $\mathrm{Cdc42^{GTP}}$ in their active forms, and $\bar{R}$ and $\bar{C}$ the concentrations of $\mathrm{RhoA^{GDP}}$ and $\mathrm{Cdc42^{GDP}}$ in their inactive forms. We can add the three conservation equations for the concentrations:

- $R_{\text{tot}} = \bar{R} + R + GR$
- $G_{\text{tot}} = G + GR$
- $C_{\text{tot}} = \bar{C} + C$

which leads to the following system of equations:

$$\begin{cases} \dfrac{dR}{dt} = k_1 . G_{\text{tot}}(R_{\text{tot}} - R - GR) - k_2 R - \dfrac{dGR}{dt} \\ \dfrac{dGR}{dt} = k_3 R(G_{\text{tot}} - GR) - k_4 GR \\ \dfrac{dC}{dt} = k_5 G_{\text{tot}}(C_{\text{tot}} - C) - k_6 C \end{cases}$$

We added few hypotheses, to reduce the number of unknown parameters:

- many GAPs are common to Cdc42 and RhoA (Müller et al., 2020), we considered their deactivation rates to be similar: $k_6 \approx k_2$.
- GTPases activated with strong stimuli -giving rise to strong phenotypic changes- lead to only 5% of the proteins in a GTP-state, both for RhoA and Cdc42 (Benard et al., 1999; Pertz, 2010; Ren et al., 1999). Therefore, we assumed that $C << C_{\text{tot}}$ and $R << R_{\text{tot}}$, even at high concentrations of optoPRG. We also considered that a few proportions of both optoPRG and $\text{RhoA}^{\text{GTP}}$ were in complex, so that $GR << R_{\text{tot}}$ and $GR << G_{\text{tot}}$.
- As we saw that the inhibition of RhoA by the PH domain of optoPRG was faster than the activation of RhoA, we considered that the reaction (*b*) was constantly at equilibrium. Thus, the complex *GR* can be expressed as a function of $G_{\text{tot}}$ and $R$ (which will both evolve in time), as the following: $GR(t) = \frac{G_{\text{tot}}(t)R(t)}{K_b}$

With these hypotheses, before optogenetic activation, we have the following equilibria:

$$\begin{cases} R_{\text{eq}} = \dfrac{G_{\text{eq}} R_{\text{tot}}}{K_a} \\ GR_{\text{eq}} = \dfrac{G_{\text{eq}} R_{\text{eq}}}{K_b} \\ C_{\text{eq}} = \dfrac{G_{\text{eq}} C_{\text{tot}}}{K_c} \end{cases}$$

with $G_{\text{tot}}(t < 0) = G_{\text{eq}}$.

#### Modeling RhoA dynamics

Introducing the new variables $r = R/R_{\text{eq}}$, $c = C/C_{\text{eq}}$, $gr = GR/GR_{\text{eq}}$ and $g = G_{\text{tot}}/G_{\text{eq}}$ leads to the much simpler and independent equations:

$$\begin{cases} \dfrac{dr}{dt} = k_2(g - r) - \dfrac{GR_{\text{eq}}}{R_{\text{eq}}} . \dfrac{dgr}{dt} \\ \dfrac{dc}{dt} = k_2(g - c) \end{cases}$$

Expressing $\frac{dgr}{dt}$ as a function of r and $\frac{dr}{dt}$, we obtain the following equation for the evolution of *r*:

$$\frac{dr}{dt} = \frac{1}{1 + \frac{G_{\text{eq}} * g}{K_b}} \left( k_2(g - r) - \frac{G_{\text{eq}} * r}{K_b} \frac{dg}{dt} \right)$$

Which contains only two unknown parameters : $k_2$ and $G_{\text{eq}}/K_b$. This second parameter will be the free parameter changing from cell to cell, due to the difference in optoPRG expression.

To evaluate $k_2$, we considered the measurements made with the biosensor of RhoA activity. This biosensor has its own intrinsic dynamic associated to a given $k_{\text{on}}$ and a $k_{\text{off}}$. Considering that only a small portion is bound to $RhoA^{GTP}$ at any moment, the dynamic of the biosensor will follow the one of $RhoA^{GTP}$ with a delay, given by $k_{\text{off}}$ in the following equation: $\frac{db}{dt} = k_{\text{off}}(r - b)$ where $b$ is the relative changes of the biosensor intensity.

$k_2$ and $k_{\text{off}}$ can be estimated independently. When the PH domain is recruited alone, if we consider that the endogenous activities of GEFs and GAPs are slow compared to the binding of the PH to $RhoA^{GTP}$, we obtain $1 - rt \propto 1 - p$ where $p$ is the relative amount of PH domain. Thus, $k_{\text{off}}$ can be fitted independently of the $k_2$, and we found that $k_{\text{off}} = 0.08 \pm 0.4 s^{-1}$ (Figure 1).

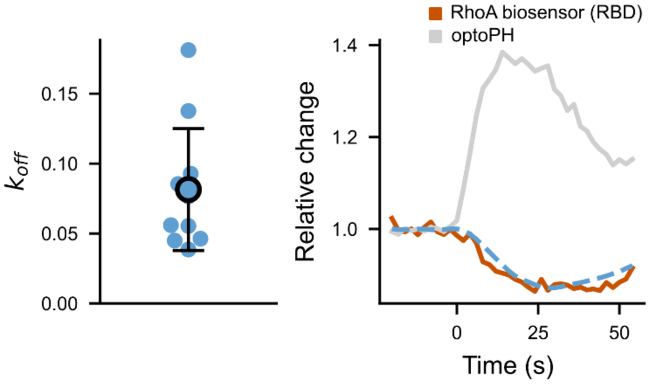


**Figure 1: Fit of the biosensor $k_{\text{off}}$.** On the left, $k_{\text{off}}$ values fitted from different cells (N=3 cells, with repeated measurements). On the right, an example of a fitted RBD curve (fit in blue, data in brown), plotted together with the normalized optoPH recruitment (gray curve).

We can than estimate $k_2$ by looking at cells with a very low concentration of optoPRG. In these cells, we expect the formation of the complex to be negligible compared to the activation of RhoA by the PH domain. Thus, the evolution of r will be given by the simple equation:

$$\frac{dr}{dt} = k_2(g - r)$$

Taking the mean value of the $k_{\text{off}}$ found above, we can fit three different dynamics in three different cells with low concentration, as shown in Figure 2, which gives $k_2 = 0.014 \pm 0.003 s^{-1}$. As seen here, this very simple model seems to well represent the data that we get by inducing pulses of RhoA activity at low concentration.

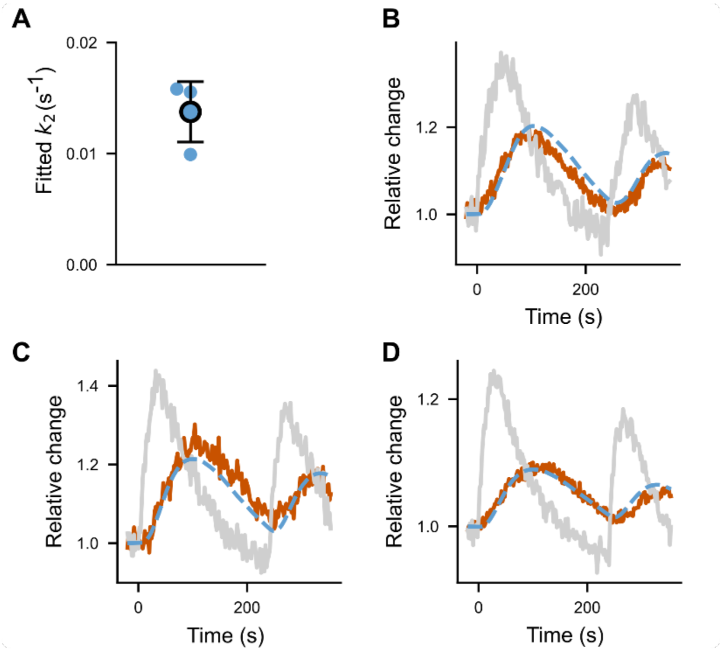


**Figure 2: Fit of $k_2$.** (A) $k_{\text{off}}$ fitted for different cells (N=3 cells). (B), (C), (D) the three fitted curves (fit in dotted blue, data in brown), plotted together with the normalized optoPH recruitment (gray curve).

As stated above, we are left with one free parameter, $G_{\text{eq}}/K_b$, for the equation describing the evolution of RhoA$^{\text{GTP}}$, which is the parameter changing from cell to cell, due the different intensities of transfection.

Taking the two mean values of $k_{\text{off}}$ and $k_2$ fitted above, we could reproduce the very different dynamics we had observed with the RBD biosensor by fitting only the ratio $G_{\text{eq}}/K_b$, as shown in Figure 3A of the main manuscript. Indeed, we recovered the behavior of low transfected cells (low $G_{\text{eq}}/K_b$) where only the activation of RhoA is present, the behavior of highly transfected cells (high $G_{\text{eq}}/K_b$) where the recruitment of optoPRG leads to a inhibition of RhoA, and the behavior of cells with medium transfection, where complex dynamics with both inhibition and activation could be well fitted.

#### Adding Cdc42 to model the double phenotype

Now, let us consider RhoA and Cdc42 responses during five minutes with simulated optoPRG inputs comparable to the one of the experiments. For $G_{\text{eq}}/K_b$, we choose values ranging between the extreme boundaries found by fitting the different curves previously.

As said in the main manuscript, the activity of RhoA will be in competition with the activity of Cdc42, that is also activated by optoPRG. The resulting phenotype will be a retraction if the absolute increase of RhoA$^{\text{GTP}}$ is superior to the absolute increase of Cdc42$^{\text{GTP}}$, with an unknown factor $\alpha$ that represents the efficiency of both GTPases to produce their specific phenotype. As we saw that the resulting phenotype was determined two minutes after the first activation, we choose this time point as the one where the other cellular processes and feedbacks take over. Thus, we compared the whole activity of RhoA and Cdc42 during the two first minutes after activation. This gives the following conditions:

$$\begin{cases} \int_0^2 (R - R_{\text{eq}})dt > \alpha \int_0^2 (C - C_{\text{eq}})dt \Rightarrow \text{retraction} \\ \int_0^2 (R - R_{\text{eq}})dt < \alpha \int_0^2 (C - C_{\text{eq}})dt \Rightarrow \text{protrusion} \end{cases}$$

We can define $\alpha' = \alpha \frac{C_{\text{tot}} K_a}{R_{\text{tot}} K_c}$ to have the equivalent condition:

$$\begin{cases} \gamma > 0 \Longrightarrow \text{retraction} \\ \gamma < 0 \Longrightarrow \text{protrusion} \end{cases}$$

where $\gamma = \int_0^2 (r-1)\,dt - \alpha' \int_0^2 (c-1)\,dt$

At low concentrations, *r* and *c* have the same dynamic, while the phenotype is always a retracting one. Therefore, we expect $\alpha'$ to be lower than one.

For the standard type of activation, we saw that the switch between retracting and protruding phenotype was happening for $G_{\text{eq}}/K_b \approx 0.5$. Thus, we choose $\alpha' = 0.24$ so that $\gamma\left(G_{\text{eq}}/K_b = 0.5\right) = 0$. We also added a 'gray zone' around 0, where the cell would not do a mixed phenotype because activities of RhoA and Cdc42 are of the same magnitude. This gray zone will be materialized by an arbitrary value, such that $|\ \gamma\ | < 3 \Longrightarrow mixed\ phenotype$.

#### Modeling optogenetic pulses

To model the pulses of optoPRG recruitments, we constructed a simple model of equilibrium between iLID at the membrane and SspB in the cytosol. We considered that with one pulse of blue light, the quantity of high affinity iLID increases with a factor *f*, and then returns to equilibrium with a half-life of $\tau_{1/2} = 20s$ (Guntas et al., 2015). We knew that PRG has a certain affinity for the membrane, we thus added a second timescale $1/k_{\text{off}}$ to take it into account.

With this, and the hypothesis that SspB is in excess, we obtain the following equation for one pulse of activation: $\frac{dg}{dt} = k_{\text{off}}(1 + (f-1) * e^{-\frac{t}{\tau_{1/2}}} - g)$. This equation can be solved analytically, and f and $k_{off}$ can be fitted to a standard pulse of activation, as seen in Figure 3.

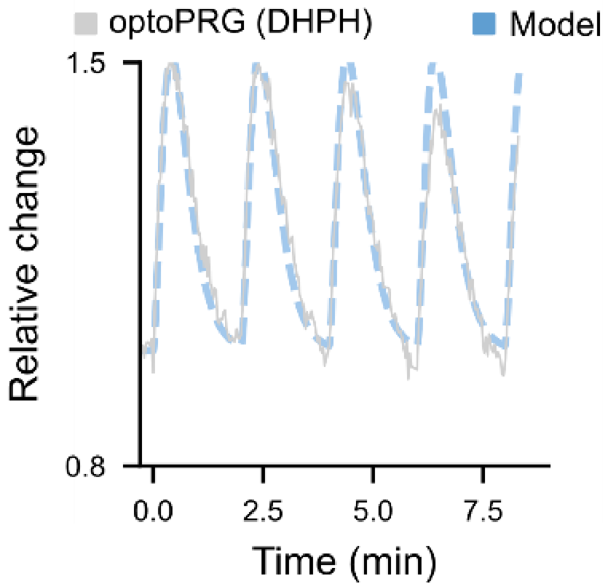


**Figure 3: fit of *f* and $k_{\text{off}}$ for pulses of optogenetic recruitment**. Pulses of light every 120 seconds lead to highly reproducible optogenetic recruitments that can be fitted with the model previously described. The values here are $k_{\text{off}} = 1/15$ s$^{-1}$ and $f = 1.5$, which are the chosen values for the model.

We used these values of $k_{\text{off}} = 1/15$ s$^{-1}$ and $f = 1.5$ for all simulations shown in the Figure 6 of the main manuscript.
